# The LAST mile: Evaluating genetic biocontrol as a supplemental tool for eradicating invasive rodents on islands

**DOI:** 10.1101/2025.11.20.688920

**Authors:** Aysegul Birand, Bruce A Hay, Matthew Combs, Luke Gierus, Katherine E Horak, Kevin P Oh, Louise J Robertson, Paul Q Thomas, Ana M. Velasquez-Escobar, David J Will, Maciej Maselko, Antoinette J Piaggio

## Abstract

Invasive rodents cause severe ecosystem degradation on islands and can be challenging to eradicate. Current best-practices rely heavily on the application of toxic oral baits and have led to successful eradications and remarkable recoveries of native flora and fauna. Yet this single method is not universally applicable. Reliance on a single method limits further eradication success in situations where toxicants cannot be used or remnant populations persist after application. Genetic biocontrol offers a suite of new solutions to potentially improve outcomes in the critical “last mile” of eradication. These approaches involve the release of genetically modified individuals of the target species to reduce population fitness over time. These include self-limiting approaches which require multiple releases and self-sustaining mechanisms (i.e., select gene-drive systems) which could theoretically collapse populations after a single release. While gene drive systems have received significant attention, their development in vertebrates remains technically challenging, and their ecological and regulatory implications are still in active debate. In contrast, some non-drive genetic biocontrol approaches, such as Y-linked editors, fsRIDL, and Gravid Lethal, offer self-limiting alternatives that may be more immediately deployable. These approaches could be used to supplement toxicant-based methods and may also have reduced environmental risks and regulatory barriers. To evaluate the potential of these tools, we developed an individual-based model simulating the eradication of house mice (*Mus musculus*) on a 125ha island with an initial population of 11,000 individuals. We tested various combinations of genetic biocontrol release and effort strategies to understand tradeoffs between required effort and uncertainty for achieving successful eradication. Under certain effort strategies, we found that the Gravid Lethal approach performed the best, achieving eradication in less than 2.3 years with intensive release effort and monitoring. Our findings suggest that integrating non-drive genetic biocontrol into adaptive management frameworks could enhance the effectiveness and feasibility of rodent eradication programs. These tools are not replacements for toxicants but may serve as critical supplements—particularly in the “last mile” of eradication. Further stakeholder engagement is needed to assess the ecological, ethical, and logistical dimensions of deploying these technologies in real-world settings.

## Introduction

Removal of invasive rodents, specifically rats (*Rattus* spp.) and house mice (*Mus musculus*), from islands can lead to significant biodiversity recovery and positive trends in overall ecosystem recovery. This recovery contributes to improved land-sea connectivity, carbon sequestration, nutrient cycling, and island community ecosystem health (Jones et al. 2016; de Wit et al. 2020; Sandin et al. 2022; Graham at al. 2024; Honzák et al. 2024). Over the last 100 years, invasive mammal eradications on islands have increased in complexity and been implemented by a majority of nations with islands (typically small and uninhabited ones), having a success rate of nearly 88% (Spatz et al. 2022). As such, invasive mammal eradications are recognized as one of the most effective conservation practices to prevent extinctions and halt environmental degradation (Langhammer et al. 2024).

Despite these successes, rodent eradication efforts remain heavily reliant on a single tool: the large-scale distribution of toxicants. While toxicants have a long and well-documented history of use in rodent control (Howald et al. 2007; Spatz et al. 2022), and best practices for their application are well established (e.g., Thomas et al. 2017; Broome et al. 2017, 2019; Keitt et al. 2015), their use is not universally feasible. Environmental constraints, non-target species risks, regulatory barriers, and social acceptability can all limit toxicant application. Moreover, eradication attempts do not always succeed on the first try. Remnant populations may persist due to bait aversion, sublethal exposure, or inaccessible terrain, leading to recolonization and failure to achieve long-term conservation outcomes (Holmes et al. 2015; Samaniego et al. 2021).

These challenges highlight the need for adaptive management—an iterative process of implementing, monitoring, and adjusting interventions in response to ecological feedback. While adaptive management strategies that rely on a successive series of labor and time-intensive removal methods to achieve complete removal are commonly used when targeting other invasive species on islands, rodent eradication programs have largely relied on static, single-method strategies to achieve the overarching set of eradication principles (Cromarty et al. 2002; Parkes 1993). As managers increasingly target larger, more complex islands—including those with human communities—this rigidity limits the effectiveness and scalability of eradication efforts (Harper et al. 2020; Oppel et al. 2011; Glen et al. 2013; Capizzi et al. 2024 Horn et al. 2022). Ambitious landscape-scale initiatives such as Predator Free 2050 (PF2050) in New Zealand exemplify the growing demand for innovative methods and adaptive management. PF2050 aims to sequentially eliminate multiple invasive mammal species from management zones, including urban areas, using natural features as reinvasion barriers (Parkes et al. 2017; Bell et al. 2019; Patterson et al. 2024). As the size and complexity of target ecosystems increase, so too does the urgency for tool diversification that can support adaptive, multi-phase eradication strategies. Thus, a range of novel methods should be explored and developed, particularly those that can be used in combination with proven and impactful toxicant interventions (Combs et al. 2023; Campbell et al. 2015; Murphy et al. 2019; Innes et al. 2024).

Innovations, such as the genetic biocontrol strategies discussed below, are unlikely to replace current best practices such as toxicant application but may provide valuable synergies when combined with them or other future methods. Most island rodent eradication efforts occur via aerial application of toxicants by helicopter or drone, or through bait stations or hand spreading (Spatz et al. 2022). Successful toxicant-based rodent eradications sometimes require multiple attempts (MacKay et al. 2007; Holmes et al. 2015; Samaniego et al. 2021; Springer et al. 2024). This can occur because of avoidance of baits, bait interference from invertebrates, and insufficient geographic and topographic coverage (MacKay et al. 2007; Holmes et al. 2015; Samaniego et al. 2021; Kappes et al. 2019). Following an initial application, greater than 90% of the original population may be removed, but remnant survivors lead to population rebound over weeks to months to pre-treatment numbers (Hein et al. 2015). Upon reviewing these unsuccessful efforts (Springer et al. 2024), there is often not a single defining reason that the implemented operational strategies did not achieve success. Instead, managers are left with several nuanced research questions that can only be addressed through years of research or high-risk full-scale operational experimentation to test the efficacy of minor adjustments in strategy.

Current best practices say that if the eradication goals cannot be met with a high degree of confidence through the use of a toxicant, then eradication should be adaptively managed using a series of progressively more resource-intensive methods to achieve complete elimination of the population. However, for rodent eradications, detecting and eliminating this “last mile” remnant post-toxicant treatment population remains an enormous challenge (Kappes et al. 2019; Davis et al. 2023). This is particularly true on more complex island systems where features such as difficult terrain, human habitation, and challenging climate create increased opportunities for persistence and rebound of remnant survivors. These points are highlighted in an assessment of recently unsuccessful eradication attempts on the globally important islands of Gough, Tristan da Cunha, and Midway Atoll, United States (Springer et al. 2024). However, recent advances in monitoring technology promise to help more effectively identify remnant rodents (Martinez et al. 2020; Piaggio et al 2024; Sullivan et al. 2024). These advances, in combination with the use of emerging genetic technologies for biocontrol may provide novel, complementary tools for achieving success in this last mile across island eradication efforts (Piaggio et al. 2017; Teem et al. 2020). Understanding the contexts in which these new genetic biocontrol tools can contribute, and how to optimize combinatorial deployment strategies in the context of rodent behavior and ecology remains a critical step in the effort to increase the success of island eradications (Combs et al. 2023).

Genetic biocontrol comprises a wide range of approaches that aim to suppress a target population following release of organisms with altered genomes (Teem et al. 2020). Genetic biocontrol can be broadly divided into two classes: (1.) those which bias their own inheritance, namely gene drives (Burt 2003), which can be self-sustaining or self-limiting and; (2) those that are non-driving, self-limiting methods that require repeated delivery in order to maintain the genetic element (or edits it creates) in the wild population (Teem et al. 2020). Both classes have the appeal that they leverage natural migration and mating behavior (the desire to seek out conspecifics) to bring about diffusion from the deployment location into the larger target environment -a key distinction from conventional toxicant- or trapping-based approaches.

Several synthetic drive strategies have been proposed, and in some cases implemented, for population suppression in insects (reviewed in Bier 2022; Johnson et al. 2024; Beach and Maselko 2025; Tolosana et al. 2025). These positive attributes notwithstanding, modeling suggests that even a relatively invasive gene drive (able to spread relatively rapidly from low frequency), such as the mouse *t*_CRISPR_ system could require a decade or more, if used alone, to eradicate an island population of ∼200,000 mice (Birand et al 2022; Gierus et al. 2022). Thus, while gene drives hold great potential (because the overall effort following initial deployment can be low), the timeframes involved are long and to date have proved inefficient in or non-transferable to mammals (Grunwald et al. 2019; Pfitzner et al. 2020; Weitzel et al. 2021; Bunting et al. 2024). Furthermore, their potential to impact populations beyond the designated treatment area imposes higher social, cultural, and regulatory barriers.

These facts argue for consideration of other genetic biocontrol strategies in which non gene-drive strategies that are intrinsically self-eliminating could be deployed strategically, following an initial population knockdown by toxicants, to increase the likelihood of crossing the last mile from suppression to eradication on islands in which eradication is currently challenging. Non-driving, self-limiting genetic biocontrol strategies utilize repeated restocking of individuals carrying the biocontrol element to maintain their effect on the population, such as the highly successful classical biocontrol methods, such as Sterile Insect Technique (SIT) (Knipling 1955; Schwarzländer et al. 2024). Although they require the release of more individuals than gene drives to suppress the target population, the development times may be shorter and the regulatory, social, and cultural pathway for the use of such elements more feasible.

Here we consider three transgene-based (GMO) genetic biocontrol strategies: female-specific Repressible Inducible Dominant Lethal (fsRIDL), Y-linked editors, and the Toxic Male Technique (TMT). With fsRIDL, homozygote males are released into the wild (Fu et al. 2007). These carry a tetracycline-repressible toxin (repressed during rearing) that would be otherwise expressed in female progeny (in environments in which tetracycline is not present), resulting in their death. With repeated releases, the generation over generation decrease in the number of females can lead to population suppression or extinction. A Y-linked editor consists of a site-specific DNA-modifying enzyme such as Cas9 and gRNAs, located on the Y chromosome. These cleave a target sequence on a different chromosome (an autosome or the X chromosome) during adult male spermatogenesis.When these males mate with wild-type (WT) females the sequence modifications inherited by progeny bring about dominant female-specific lethality or infertility of progeny. Because the construct acts in the adult male germline to reduce the fitness of female progeny, and the Y-chromosome is not present in these individuals, the construct is not selected against by the harm it causes. As a result, under ideal conditions (no fitness cost in carrier males), the transgene persists in the population at its introduction frequency. Thus, releases of transgene-bearing males over multiple generations progressively ratchets the population to a terminal state consisting of fertile males and infertile or dead females (Burt and Deredec 2018). Finally, the idea behind the Toxic Male Technique (TMT) is that males are engineered to produce one or more toxins in their seminal fluid. The hope is that when these are transferred to females during mating, the mated females will die (Beach and Maselko 2025).

TMT is different from the first two approaches in that it acts during the release generation. An important corollary is that TMT’s suppression ability, unlike that of other strategies (Gierus et al. 2022; Birand et al. 2022), benefits from polyandry, which is common in invasive mice. It could also be very useful for longer lived species where conventional intergenerational transmission may take decades to see an impact. Although TMT is likely to be difficult to engineer in rodents, we reasoned that a TMT – like approach (theoretical at this point) which could rapidly reduce the reproductive capacity of a target population by eliminating mated females may be effective. Here we model a version of TMT called Gravid Lethal. In this system males are engineered to carry a transgene that, when expressed by the placental or embryonic tissues of their offspring, results in the death of the gravid female. This could potentially be achieved via prenatal (embryo-specific) expression of small molecular weight toxin proteins or enzymes producing toxic secondary metabolites, with the toxin acting non-autonomously to kill the mother. A drug-dependent repressible regulatory circuit (e.g., tet-off) could be used to ensure the transgene is not expressed in lab populations.

For each of the above methods we use modeling to explore if there are genetic biocontrol release efforts and monitoring strategies that could lead to successful eradication when used following application of a toxicant that brings about an initial large decrease in population size (>90%).

The idea is that well-validated toxicants can be used to rapidly decrease population numbers, thereby providing a context in which modest size releases of non-driving, self-limiting genetically modified animals of a single sex (typically males) can be used to bring about final eradication of remnant populations. This approach potentially overcomes the problem of needing to release large numbers of the invasive species required by an unassisted self-limiting genetic biocontrol approach or releasing animals carrying a self-sustaining gene drive element. Further, it may lead to broader use applications when toxicant alone could not achieve the last mile. We recognize that eradication efforts using genetic biocontrol, specifically self-limiting based techniques, come with the need for numerous releases of genetically modified animals over long timelines, as compared with large scale aerial toxicant bait applications. However, because some island eradications efforts utilizing only toxicants fail or have other complicating factors (geography, size, a mix of wildlands and human occupation, etc) we believe it is important to ask if strategies such as genetic biocontrol, which take advantage of the target species desire to seek out conspecifics for mating, can broaden the possibility for successful eradications to island systems that are currently infeasible. We propose that this approach (LAST; LAst mile Suppression Technology) could fit into the broader pest control paradigm of using a series of methods or tools and adaptive management to achieve eradication.

While the LAST approaches are attractive in principle, to understand contexts in which they could realistically contribute, it is critical to know how many genetic biocontrol individuals would need to be released, how often releases would occur, and what release, monitoring, and management strategies would be most successful. Modeling genetic biocontrol supplementation to toxicant application can allow us to better understand these parameters. With the goal of characterizing parameters and application effort strategies necessary for successful eradication, we use the stochastic, spatially explicit individual-based modeling framework presented in Birand *et al*. (2022) to investigate the efficacy of selected genetic biocontrol methods, following an initial toxicant-mediated reduction in population size, for the eradication of invasive mice, as a rodent model, on islands.

## Methods

### The model

The model is a patch-based stochastic model where individuals occupy a rectangular array of patches, and each patch holds multiple individuals. However, individuals are not restricted to a single patch but can utilize multiple patches even within a single breeding cycle. Each breeding cycle is considered a time-step in the model, there are multiple breeding cycles in a year, and generations are overlapping. The following steps occur in each breeding cycle: 1) mate search, 2) mating, 3) reproduction, 4) natal dispersal, 5) survival of adults, and 6) breeding dispersal (for more details, see ‘The model” Appendix S1).

#### Control strategies

Two control strategies were employed: *lethal control* (toxicant) followed by *genetic biocontrol* (non-driving). Lethal control was simulated as a range of proportional culling strategies operating on the population. When control was implemented, survival probabilities of all individuals were reduced by multiplier 1 -*c*_s_ (reduction in survival with lethal control) per breeding cycle. Subsequently, genetic biocontrol was employed with a release of transgenic males of each of the LAST approaches under varying release strategies with varying effort (see below).

Three genetic biocontrol strategies (LAST) were investigated: fsRIDL, Y-linked editor, and Gravid Lethal. With fsRIDL, WT females mate with transgenic males and the female offspring are transgene-bearing and suffer mortality, whereas with the Y-linked editor, the female offspring of the WT females mated with transgenic males are sterile. With the use of Gravid Lethal, the WT females mated with transgenic males suffer mortality without producing any offspring due to expression of a toxic transgene in prenatal development that causes death of the mother.

#### Release strategies for genetic biocontrol

We modeled three release strategies for each genetic biocontrol strategy: *broad*, *tactical*, and *tactical when-detected*. Under *broad* release, transgenic mice were released to all release sites (*n*_p_). With the *tactical* release strategy, there was monitoring at initial release sites for the presence of individual mice (during the entire release duration, *n*_i_). Transgenic mice were then released only to the sites where mice were detected in the previous breeding cycle (Fig. 1). In the final release strategy, *tactical when-detected*, we assumed that transgenic mice could be differentiated from WT mice during environmental monitoring for DNA (Piaggio et al. 2024). Under this release strategy, releases occurred only if WT individuals were detected, thus reducing the overall release effort and cost to releasing introduction of transgenic mice when possible. Note that each release strategy implicitly assumes variable monitoring efforts, i.e. no monitoring effort with the *broad* release, medium monitoring effort the *tactical release*, and high monitoring effort with the *tactical when-detected* release.

**Figure 1.**
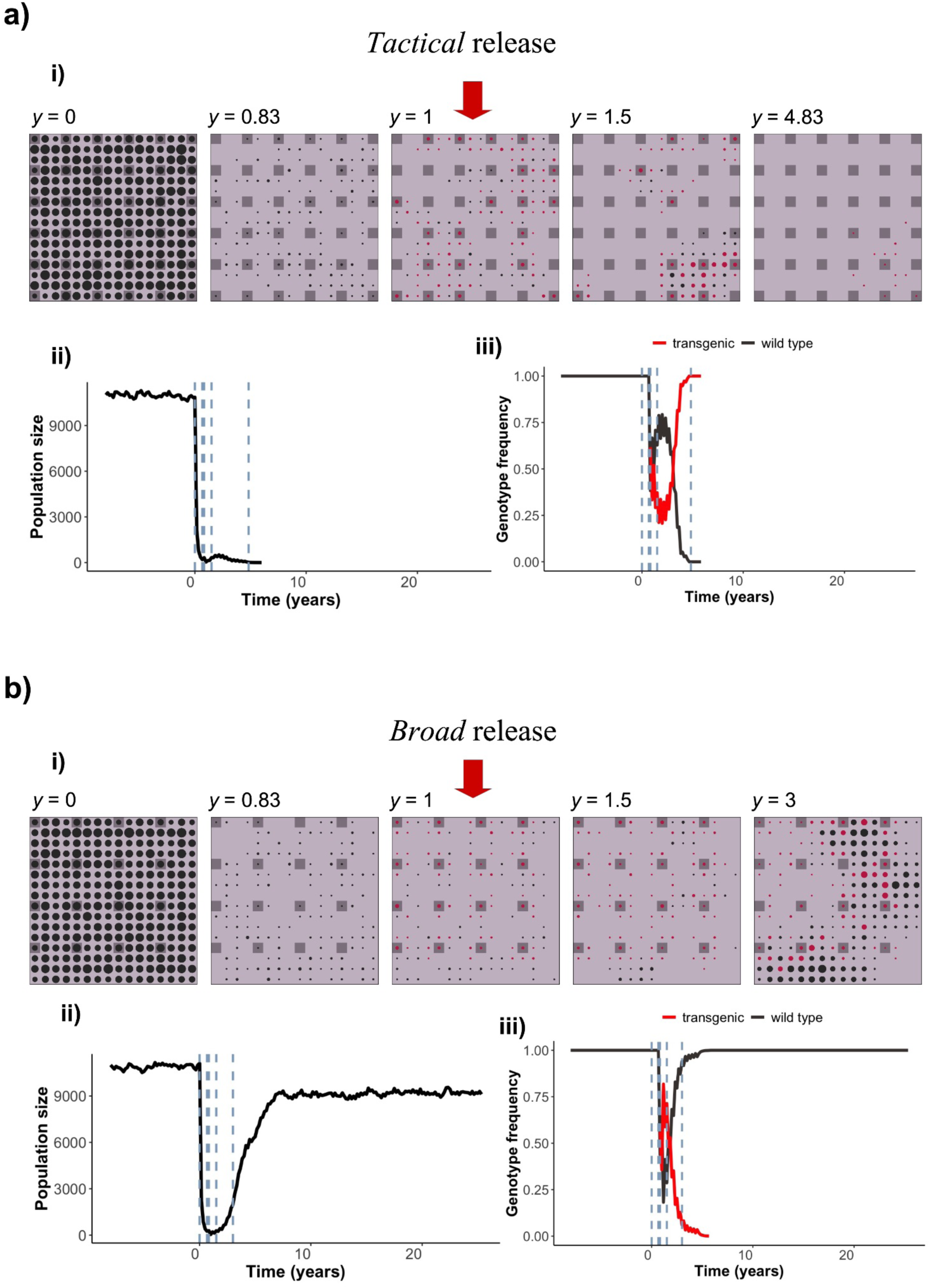
Progression of two sample simulations through time using the Gravid Lethal biocontrol strategy with *tactical* (**a**) and *broad* (**b**) release. Each square plot in (**i**) represents the hypothetical island (as an array of 16 x 16 square patches) with mouse populations through time starting at the end of the eight-year burn in period (*y* = 0). Populations in patches are represented as circles, the size of which is proportional to the mouse population size in that patch. The black circles represent populations with wild type individuals only; red circles represent populations with one or more transgenic individuals. Dark gray squares are patches where transgenic individuals are released, the same patches are also monitored under *tactical* release in ***a***. Empty patches are light gray. The introduction of transgenic individuals started at *y* = 1, after toxicant application l knocked down the population. Released genetic biocontrol individuals can disperse from the release sites before joining the mating pool. **ii)** Population size on the island through time. The vertical dashed lines correspond to the times of the simulation snapshots in subplot ***i***. **iii)** The genotype frequencies of wild type and transgenic individuals through time. Eradication was successful in 6 years in ***a***, and failed in ***b***. The population size after toxicant application and before genetic biocontrol releases was 39 in ***a***, and 42 in ***b***. Transgenic mice were introduced to 36 patches in ***a***, and 16 patches in ***b***. (Other parameters are: *n*_i_ = 12, *N*_i_ = 10, *c*_s_ = 0.20, *D* = 1, *p*_r_ = 0.8, *p*_m_ = 0.8; videos of these simulations are available at https://github.com/abirand/GeneticBiocontrol)

#### Release effort

For each release strategy presented above, we varied the overall release effort by changing the spatial effort (*n*_p_, the number of release sites distributed systematically across the island), temporal effort (number of breeding cycles when releases occurred, *n*_i_), and the number of individuals released per patch (*N*_i_) the three dimensional ‘effort cube’ (Fig. 2) demonstrates the release combinations investigated in this study. The overall release effort is the total number of transgenic males released, which is *N*_T_ = *N*_i_*n*_p_*n*_i_. Note that this definition of effort does not include costs associated with monitoring. In order to accommodate the potential logistical challenges that might occur, we assumed that the releases occurred every other breeding cycle, i.e *n*_i_ = 6 corresponds to 2 years since we assume that there are 6 breeding cycles (*n*_c_) in a year. After being released, for all strategies, transgenic males randomly choose a patch within distance *D* from their introduction sites and join the pool of available males for mating. Females (with probability *p*_r_) choose males randomly among all the available males within the patch, which implies that not all females mate, and some transgenic males may not be chosen for mating. We also assume there is no bias towards mating with WT males.

**Figure 2.**
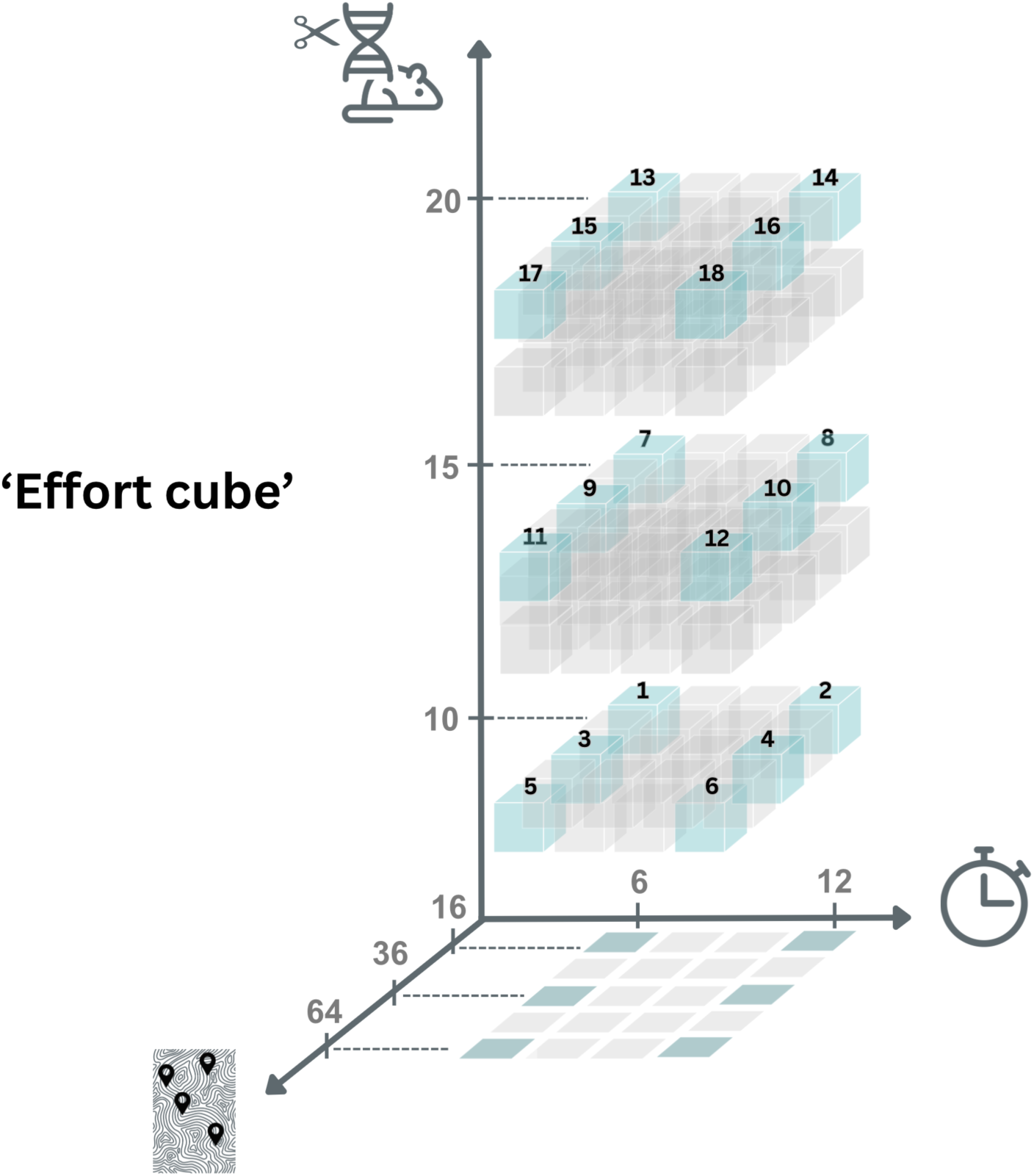
The ‘effort cube’ represents release strategies with varying spatial (*n*_p_, represented by map graphic), temporal (*n*_i_, represented by clock graphic), and per-site introduction (*N*_i_, represented by mouse/DNA/scissors graphic) efforts. Numbers on the axes represent the values used in the simulations (see Appendix S1, Table S1), whereas the numbers on the cubes represent combinations of effort modeled under each release strategy: i.e. #1 represents low spatial effort (*n*_p_ = 16 patches), low temporal effort (*n*_i_ = 6 times) and low per-site introduction effort (*N*_i_ = 10 transgenic males); whereas #18 represents high effort for all three (64, 12, and 20, respectively). The ‘shadow’ presented in the graph is used to represent the data in Figure 3.

#### Parameters

To reduce the parameter space for efficient simulations, we selected mate search and dispersal distances (*D*), levels of polyandrous mating (*p*_m_), and proportion of females mating in the population (*p*_r_) as life history parameters with potentially the highest impact on the efficacy of genetic biocontrol (Birand et al. 2022; Gierus et al. 2022) and investigated those in detail. All parameters related to life history traits, control strategies, and release efforts are presented (Appendix S1, Table S1). Since we are interested in the ability of genetic biocontrol to lead to successful eradications upon the failure of toxicant application to achieve complete population eradication, we chose values for *c*_s_ accordingly and used the simulations in which there were still individuals left on the island (Appendix S1, Figure S1). All release effort combinations with three release strategies for each genetic biocontrol strategy and variable life history traits resulted in 1296 parameter combinations, and we ran 30 simulations for each combination (116,640 simulations in total). Release strategies that vary the combinations of these efforts (spatial, temporal, and introduction size; Fig. 2) require different degrees of overall effort and represent trade-offs that are likely to be important in different environmental and management contexts when toxicant application alone is not successful. Furthermore, we performed a global sensitivity analysis to investigate the relative influence of these parameters on the probability of successful eradication (for more details, see ‘The model’ and Appendix S1, Table S1). The effects of life history parameters that are not included in the sensitivity analysis were investigated in detail for self-sustaining genetic biocontrol strategies previously (Birand et al. 2022; Gierus et al. 2022).

#### Initial conditions

We assumed that the hypothetical island was 16 x 16 = 256 patches in the model. Each patch corresponds to a 70m x 70m space on a hypothetical island of approximately 125 hectares (ha). The maximum dispersal distances *D* = [1,3] in the model corresponds to 140 – 280 m in the wild. The probability of moving these distances increases (see step 4 in the model, Appendix S1) when the population densities are low; individuals are more likely to remain in their patches at densities close to carrying capacity. We assumed that the population size was ∼11,000 before toxicant application. We ran simulations for 200 breeding cycles (33.3 years), which had an initial burn-in period of 8 years where no control was applied, and a subsequent 1-year period during which lethal control was simulated as a range of proportional culling strategies (*c*_s_ = 0.15, 0.2) operating on the population. Following the toxicant application, genetic biocontrol males were released (for sample simulations demonstrating release strategies, see Fig. 1). We labeled simulation outcomes as unsuccessful if eradication did not occur within the number of breeding cycles simulated. The model is coded using C programming language and is available at GitHub repository https://github.com/abirand/GeneticBiocontrol.

## Results

All three LAST strategies (fsRIDL, Y-linked editor, and Gravid Lethal) caused permanent eradication with some of the high effort release combinations (Fig. 3a and Appendix S1, Table S2, Figs. S2, S3), but other combinations failed. Among the genetic biocontrol strategies examined, overall Gravid Lethal outperformed the other LAST strategies, with higher eradication probabilities for more release effort combinations, particularly when the spatial and/or temporal effort were not high (Fig. 3). fsRIDL was the worst performing genetic biocontrol strategy out of the three (Appendix S1, Fig. S2), potentially because the reduced local population size due to female offspring mortality resulted in higher fertility rates (due to density-dependent fertility, see Step 3 in the Model, Appendix S1), which increased the possibility of producing WT offspring. Even though both fsRIDL and Y-linked editor were less effective compared to Gravid Lethal, release efforts affected the probabilities of eradication. Further, the expected times to eradication and the total number of individuals required in releases that resulted in eradications were qualitatively very similar for all genetic biocontrol strategies (compare Fig. 3 with Figs. S2 and S3, also Tables S2-S3 in Appendix S1). There were some effort combinations where the Y-linked editor outperformed Gravid Lethal (Appendix S1, Fig. S3). The results for the rest of this section are based on only Gravid Lethal; however, the general trends hold for the other genetic control strategies as well.

**Figure 3.**
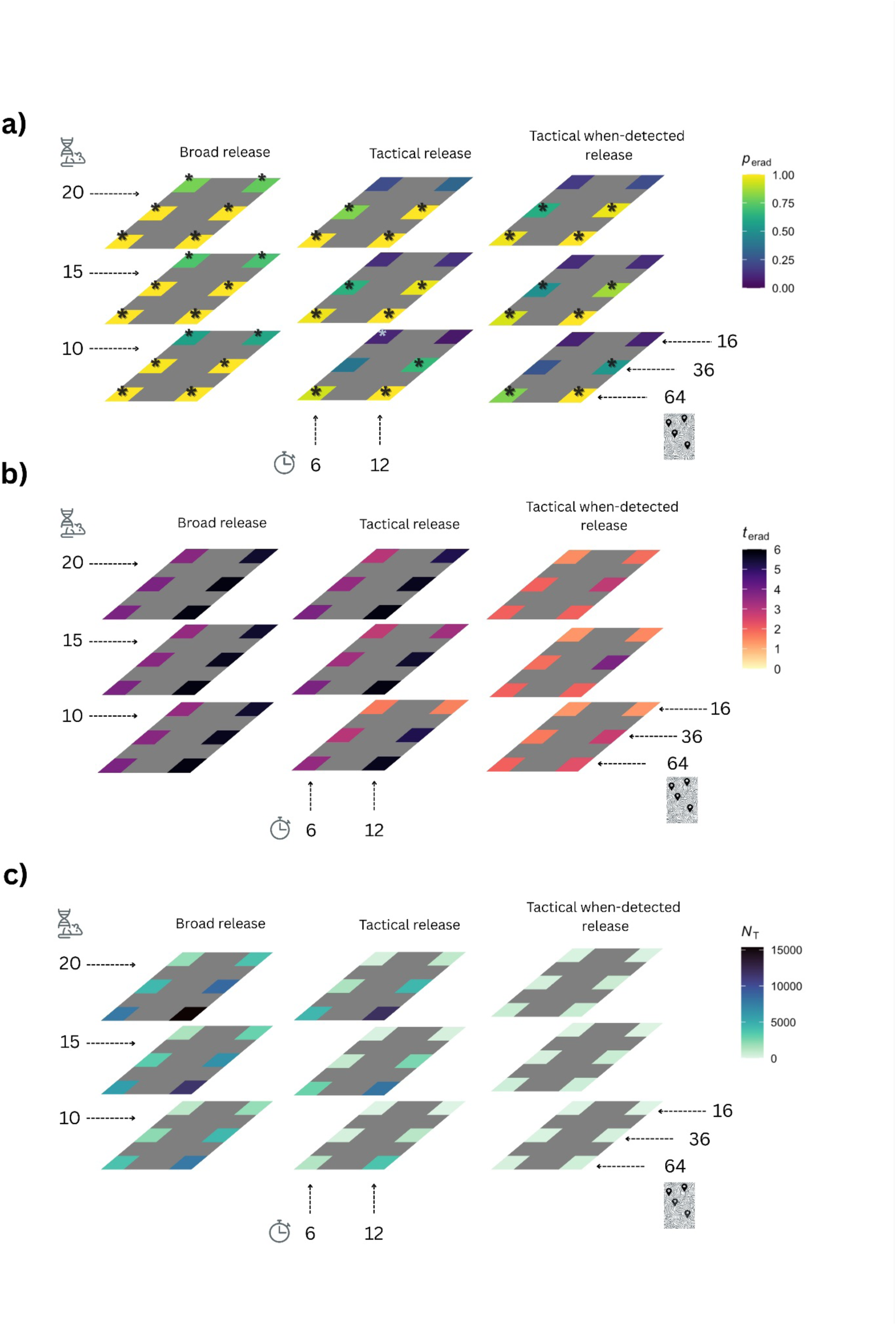
Effort cube for Gravid Lethal genetic biocontrol **a)**. The probability of eradication for each release effort combination (#1 to #18 in Fig. 1) under *broad*, *tactical*, and *tactical when-detected* release strategies. Note that the mouse/DNA/scissors graphic represents *N*_i_, clock graphic represents *n*_i_, and map graphic represents *n*_p_. The release efforts where Gravid Lethal outperformed other genetic biocontrol strategies are displayed with ‘*’. **b)** The median time to eradication (in years) for simulations when eradications were successful. **c)** The median number of transgenic individuals introduced for each effort combination. Note that the darker shades in panels **a-c** represent less desirable outcomes with either low eradication probabilities, longer times to eradication, or very high release efforts (in terms of total number of genetic biocontrol individuals introduced). The raw data are provided in Tables S2-S4, for all genetic control strategies. (Based on 720 simulations for each combination with variable rates of mate-search and dispersal distance, *D*, female probability of mating, *p*_r_, and probability of polyandry, *p*_m_).

*Broad release* (transgenic mice were released to all the predetermined release sites) outperformed *tactical release* (includes additional monitoring and subsequent releases when presence of mice were detected) with higher probabilities of eradication for more release effort combinations (Fig. 3a). Similarly, *tactical release* outperformed the *tactical when-detected release* strategy (releases occurred only if WT individuals were detected through environmental monitoring for DNA).

However, both *tactical release* strategies (with or without DNA positive detection in the environment) generally had shorter times to successful eradications (Fig. 3b). For example, for the highest overall release effort with 20 transgenic mice per patch to 64 patches, 12 times (release effort combination 18, Fig. 2) all three releases were successful (with probability of eradication = 1, Appendix S1, Table S3), the median expected times to eradication were 5.83 for both *broad* and *tactical release* strategies, and was 2 years with the *tactical when-detected release* strategy. This means that eradication was often achieved before the total number of releases were completed (e.g. the expected median time for successful eradications with release effort combinations 6, 12, and 18 were 2 -2.33 years, which correspond to temporal effort of *n*_i_ = 6,7 releases; see Appendix S1, Table S3). Overall, increasing release effort on all three dimensions resulted in higher eradication success (Fig. 3a).

In cases where eradication was successful, the total number of transgenic mice introduced was also much lower with *tactical release*, which was further reduced with the *tactical when-detected release* strategy. For example, when 10 transgenic mice per patch were released to 64 patches under the *broad release* strategy for 12 release times (release effort combination #6, Fig. 2) there were a total of 7680 transgenic mice released. However, only 3730 and 410 transgenic mice were released under the *tactical* and *tactical when-detected* release strategies, respectively (for all release effort combinations using three genetic biocontrol strategies, see Appendix S1, Table S4).

Spatial effort (*n*_p_, number of places where transgenic mice were introduced) had a strong impact on eradication probabilities for all release strategies (also see Appendix S1, Figure S4, for the relative influence of *n*_p_ on the probability of eradication). Low spatial effort (*n*_p_ = 16, release combinations 1, 2, 7, 8, 13, and 14, Fig. 2) always resulted in low eradication probabilities. Increasing temporal effort (*n*_i_, number of breeding cycles when releases occurred), and per-site introduction effort (*N*_i_, number of mice introduced per site) when spatial effort was low, did not increase the eradication probabilities further, even with the *broad release* strategy (Fig. 3a). With moderate spatial effort (*n*_p_ = 36), increasing temporal effort increased eradication probabilities (e.g. compare release strategies 3 and 4, Fig. 2). If both the spatial and the temporal release efforts were not extensive due to time and spatial constraints, increasing per-site introduction effort increased eradication probabilities (e.g. compare release strategies 3 and 9, Fig. 2).

All results presented here were pooled across simulations with variable rates for some of the most critical life history parameters on dispersal and mating that have been shown to impact the success of self-sustaining murine gene drives (Appendix S1, Table S4; Birand et al. 2022; Gierus et al. 2022). Therefore, the results presented here have included some uncertainty related to these parameters and could be considered as ‘general’ parameters for mice. Sensitivity analysis results (Appendix S1, Fig. S4) showed that the relative influence of these life history parameters on probability of eradication is low compared to the relative influence of those related to release strategies, and the effectiveness of toxicant application prior to the introduction of genetic biocontrol. Therefore, our modeling results suggest that even when the exact values for some of the life history parameters are unknown, it is possible to achieve eradication with the LAST approach when toxicant application would have failed.

## Discussion

Despite successful eradications with toxicants, their limitations in certain contexts underscore the need for supplementary tools. Incomplete eradications—due to bait aversion, inaccessible terrain, or sublethal exposure—can leave remnant populations that repopulate and continue to degrade native ecosystems (Holmes et al. 2015). The field of genetic biocontrol holds promise (Teem et al. 2020) for providing new tools. These could be used in an adaptive management context to supplement toxicant applications as a LAST approach and thus increase the likelihood of successful rodent eradication in environments previously deemed intractable.

The results of our modeling of a LAST approach for eradicating an invasive mouse island population shows that the Gravid Lethal strategy is promising as a supplementary tool for contexts in which toxicants may not achieve 100% removal alone. While all three genetic biocontrol strategies could lead to successful eradications, Y-linked editor and fsRIDL generally required the highest release efforts, i.e. *broad* releases with high spatial and temporal effort. Gravid Lethal was successful for more release strategies under *broad*, *tactical* and *tactical when detected*. This can be understood because with Gravid Lethal, WT females mate with transgenic males and suffer mortality without producing any offspring. In contrast, in the other two LAST strategies eradication is achieved via mortality (fsRIDL) or infertility (Y-linked editor) of female offspring of transgenic males mating with WT female mice. This contrasts with the other strategies modeled, in which the possibility of a female mating with multiple males, including WT, provided opportunities for creation of WT progeny, which may then also mate with other WT individuals, thereby decreasing the efficacy of the strategy. Although each of the release strategies we modeled could lead to success with Gravid Lethal, they often required high spatial and temporal release efforts (to 64 patches, 12 times). However, the expected median time to eradication with the Gravid Lethal approach using the *tactical when-detected* release strategy ranged between 2 and 2.33 years (Appendix S1, Table S3), which means that eradication was achieved after 6-7 releases–before the full 12 releases were completed–which also significantly reduced the total number of individuals introduced (410 -680 individuals). It’s important to recognize that each release strategy implicitly involves different levels of monitoring effort: *broad* release assumes minimal or no monitoring, *tactical* release requires moderate monitoring, and *tactical release when-detected* demands more intensive monitoring. There is a trade-off between the effort invested in release and the effort required for monitoring. Increased monitoring effort for 2-3 years could lead to successful eradications with more modest release effort in terms of total number of individuals introduced. If monitoring is not possible, then high release efforts are needed to achieve successful eradications.

The Gravid Lethal approach differs fundamentally because it is eliminated in a single generation and leaves no genetic trace. The lack of dependence on intergenerational transmission also renders Gravid Lethal much less susceptible to transgene mutations (e.g. rearrangements) that have been shown to lead to population resistance to gene drives, particularly self-sustaining approaches (Gierus et al. 2022). Further, although not investigated in detail here, the deployment of Gravid Lethal into a relatively small (post-toxicant) population would likely provide less opportunity for selection of alleles that are resistant to the transgene effect. Continued research and development of self-limiting genetic biocontrol options for invasive species eradication may lead to development of other approaches that allow us to achieve more successes in a shorter time-period, with the need for fewer transgenic individuals. However, any approach that is based on the creation of transgenic progeny (e.g. fsRIDL and Y-linked editors) will need to work well in the face of polyandry, which is common in rodents, and acts to defeat suppression through continued creation of wildtype progeny.

As with all wildlife management tools, no single genetic biocontrol approach is optimal for all circumstances. Multiple factors need to be considered. These include the biology and ecology of the target population, the capacity for deployment at scale, and the capability for spatial and temporal control. The power of the modeling presented herein is that it demonstrates ways these efforts can be tailored to specific applications and where failure is predicted to occur if required efforts are not achievable due to practical or logistical reasons (Fig. 3). The model demonstrates that spatial effort is important, with introduction of genetic biocontrol mice to many regularly spaced sites across the island being critical for success. Spatial considerations are also important when surviving mice are detected after toxicant application. How subsequent restocking of genetic biocontrol is deployed in the face of these detections has a large influence on success.

The insights gained by the modeling in this regard highlight situations of particular concern, such as when parts of an island are inaccessible, unless other methods of delivery can be utilized (e.g., drone or plane). It will be important for managers considering Gravid Lethal or related approaches to consider each factor carefully for their specific use case. For example, with repeated restocking, the total numbers released over the entire course of control effort may exceed the initial population size on the island, and thus there must be tolerance for short-term impacts of that introduction. If a transient population increase cannot be tolerated in a specific use case, then this tool would not be viable. Further, the rearing and repeated delivery of a high number of genetic biocontrol mice may not be financially feasible for remote regions. Like other tools, the ones presented here may be useful for some use cases but not others.

Risk assessments can be informed by the factors and outcomes modeled in this study, which will allow a careful consideration of the balance between costs, logistics, and outcome uncertainty (Combs et al. 2023). The strategy of *tactical release* and *tactical when-detected*, releasing only in areas where mice have been detected, reduces the number of mice required to be released on the island considerably. However, this strategy was successful only if the monitoring effort is extensive across space and time. In addition, the *tactical when-detected strategy* presupposes the existence of detection technologies with high sensitivity and specificity, a field still at a very early stage. The detection of target species from environmental monitoring (eDNA; e.g., water and soil) provides one such approach and has become a useful monitoring tool (Piaggio 2021) but also increases the effort and cost of eradications, a fact that needs to be considered in feasibility discussions. When monitoring after application of genetic biocontrol, an eDNA assay that can differentiate between WT mice and transgenic mice (Piaggio et al. 2024) could reduce the number of genetic biocontrol mice introduced, by reducing release efforts to only sites where WT mice are detected. When this approach is implemented in the model, the total numbers of genetic biocontrol mice as well as effort dedicated to the release of biocontrol animals and time to eradication are significantly reduced.

We also note that while the other genetic biocontrol methods we explored, fsRIDL and Y-linked editor, did not have as many pathways to success as Gravid Lethal, there were some that achieved eradication (Appendix S1, Table S2, Figs. S2, S3). The effort cube (Fig. 2, Fig. 3, Appendix S1, Table S2, Figs. S2, S3) provides a way to understand the trade-offs to implementation for each supplemental biocontrol method combined with alternative application strategies and is a useful tool for managers tasked with determining plans for successful eradications.

Finally, we emphasize that the modeling we carried out represents an initial step toward predicting outcomes of various genetic biocontrol interventions. To our knowledge, this is the first modeling study to evaluate the integration of non-drive genetic biocontrol with toxicant-based knockdown in an adaptive management framework for invasive rodent eradication (LAST approach). While individual-based models can offer valuable insights, they are inherently limited by uncertainties in rodent biology and ecology—particularly when applied to specific island contexts. Key factors such as island topology, accessibility for biocontrol deployment, genetic connectivity of populations (Oh et al. 2021), as well as breeding ecology and behavioral dynamics (Moro et al. 2018; Godwin et al. 2019), must be characterized in detail to refine

## Conclusion

Given the dire impacts of invasive rodents on islands, and the well-documented positive return on investment from their removal, there is a growing demand for new methods that expand the scope of eradication and reduce the risk of failure. To date, most applications of genetic biocontrol for invasive rodent eradication have focused on gene drives or toxicant-based strategies, often framed as all-or-nothing solutions. Our results demonstrate that self-limiting genetic biocontrol LAST strategies can serve as effective supplementary tools particularly in the “last mile” of eradication, where one strategy alone may fall short. Modeling results suggest that under certain conditions, the Gravid Lethal approach can achieve eradication following a toxicant-based knockdown. While the required effort is substantial—similar to that of current toxicant-only strategies—this approach may extend eradication feasibility to more complex or constrained environments. Importantly, it offers a path forward for adaptive, multi-method eradication frameworks. We encourage continued development and phased testing of genetic biocontrol strategies beyond self-sustaining or self-limiting gene drive approaches, with particular attention to self-limiting, non-heritable approaches that may face fewer ecological and regulatory barriers. Public engagement, transparent risk assessment, and site-specific ecological studies will be essential to ensure these tools are deployed responsibly and effectively in support of island restoration (Akbari et al. 2015; National Academies of Sciences, Engineering, and Medicine 2016; Long et al. 2020).

## Appendix S1

File format .docx

Title of data **Description of the model and the sensitivity analysis with additional results** Description of data description of model details, sensitivity analysis, and further results.

## Acknowledgements

The findings and conclusions in this publication are those of the author(s) and should not be construed to represent any official USDA or U.S. Government determination or policy. This work was also supported with supercomputing resources provided by the Phoenix High Performance Computing (HPC) service at the University of Adelaide.

## Authors’ contributions

AB contribution to the design of the study, the acquisition, analysis, or interpretation of the data, the generation of figures and tables, development of the model code for simulations, and writing of the manuscript. AJP, contribution to the conception and design of the study, led the consortium of authors in the effort, interpretation of the data, and led writing of the manuscript. MM, contribution to the design of the study, concept and design of the TMT/Gravid Lethal concept, interpretation of the data, and writing of the manuscript. BAH, MC, LG, KEH, KPO, LJR, PQT, AMV-E, and DJW, contribution to the design of the study, interpretation of the data, and writing of the manuscript. All authors read and approved the final manuscript.

## Ethics approval

“Not applicable”

## Consent to participate

“Not applicable”

