## Supplementary Materials for "The LAST mile: Evaluating genetic biocontrol as a supplemental tool for eradicating invasive rodents on islands"

### Appendix S1 Description of the model and the sensitivity analysis with additional results

#### The model

Here we provide more details about the individual-based model (Birand *et al*. 2022). The model is stochastic and spatially explicit. Individuals occupy a rectangular array of patches, and each patch holds multiple individuals. Individuals can utilize multiple patches even within a single breeding cycle. Each breeding cycle is considered a time-step in the model, and there are multiple breeding cycles in a year ($n_c$). The generations are overlapping, individuals can survive multiple breeding cycles up to a maximum age ($age_m$). The following steps occur in each breeding cycle (for more details, see Birand *et al*., 2022):

*1) Mate search.* Each breeding cycle starts with mate search. A female's search for mates begins within the female's central patch and continues in the neighboring patches incrementally with distance until either a mate is found, or the maximum search distance, $D_m$ is reached. All patches of equal distance have the same probability of being chosen during the mate search. Individuals retain their central patch within a breeding cycle, even when they fail to find a mate in that cycle.

*2) Mating.* Females mate with probability $p_r$, if they find a male within their mate-search area. If multiple males are present in a female's central patch, she can mate with multiple males ($n_m$), with probability ($p_m$). Females choose males randomly, which means that some males can mate multiple times in a single breeding cycle, whereas others may not mate at all.

*3) Density-dependent reproduction.* The number of offspring from each mated female is drawn from a Poisson distribution with mean $\upsilon = b/(1 + [(b/2) - 1][N/K])$ (discrete-time Beverton-Holt model; Kot, 2001), where $b$ is the average number of offspring of females in the absence of density-dependent regulation, $N$ and $K$ are the population size and the carrying capacity in a female's central patch, respectively. Under polyandrous mating, the paternity of each offspring is determined randomly. The sex of each offspring is determined by the sex chromosomes inherited from parents. All offspring are assumed to survive and join the mating pool in the next breeding cycle since offspring mortality is incorporated in the density-dependent reproduction function, unless there is population-wide control strategy in place (see *Control strategies* in the main text).

*4) Natal dispersal.* An offspring can disperse to a new patch from its maternal patch within distance $D_n$ to establish a mate-search area before its first breeding attempt. Natal dispersal is both distance and negative density dependent (Birand *et al*., 2022), which ensures that when the population size is close to carrying capacity, the probability of dispersing long distances is low. The probability of dispersing to distance $\delta$ is calculated as: $P(\delta)=\exp[a\ \delta/D_n-cN_r/K_r]^{2;}$; where where $a$ and $c$ are dispersal and density coefficients respectively, are both assumed to be equal to 1; $D_n$ is the maximum distance that an individual could disperse; $N_r$ is the population size, and $K_r$ is the carrying capacity, where both are both calculated by summing across all the patches within distance $D_n$. The probabilities are normalized to sum to one for each dispersing individual. All patches of equal distance have the same probability of being chosen irrespective of their direction.

*5) Survival of adults.* Adult survival probability ($\omega$) is constant for each breeding cycle unless there is population wide control strategy in place (see below). We did not assume any additional fitness costs for transgenic males.

*6) Breeding dispersal.* Surviving adults can establish new mate-search areas with a new central patch within distance $D_b$. The probability of moving a distance $\delta$ is calculated as natal dispersal above (for simplicity, we assume that the maximum distances for natal dispersal, breeding dispersal, and mate-search distance parameters are physiologically constrained to be equal, and determined by $D$). The negative density-dependent dispersal function ensures that when the population size is close to carrying capacity, individuals tend to retain the same central patch for natal and breeding dispersal and form long-term stable communities. At low densities, the probability of picking a distant patch to center its new mate-search area is higher, which can be justified by an individual's imperative to move large distances to find mates when local density is nearly zero. Life history and demography parameters are based on empirical data whenever available (Supplementary Table 4; Birand *et al*., 2022).

*Sensitivity analysis.* We performed a global sensitivity analysis for each release strategy to investigate the relative influence of parameters on the probability of successful eradication using gravid lethal as genetic biocontrol. For each of the release strategies, we created 10,000 unique parameter combinations from parameter ranges given in Supplementary Table 1 using Latin hypercube sampling (randomLHS, R package; Carnell, 2020). To minimize computational effort and maximize the coverage of the parameter space, we ran a single simulation for each parameter combination (Prowse *et al.* 2016). Finally, we examined the influence of parameter inputs using Boosted Regression Tree models (BRT; R package *dismo*; Hijmans *et al*., 2011) that we fitted to the simulation outputs using the function gbm.step from the R package *dismo* (learning rate: 0.01; bag fraction: 0.75; tree complexity: 3; and 5-fold cross-validation; Elith *et al*., 2008). We used binomial error distribution for the probability of eradication. We labeled simulation outcomes as unsuccessful for the binomial error distribution if eradication did not occur within the number of breeding cycles simulated, even when the population was suppressed to a new stable level.

**Table S1.** Parameters of the model. Base values are used as default values in most of the simulations. For sensitivity analyses (SA), parameter combinations are drawn from a uniform distribution ($U$) or uniform discrete distribution ($U_d$) using Latin hypercube sampling. Base values resulted in 3,880 parameter combinations (with three release strategies and three control strategies), and we ran 30 simulations for each parameter combination (116,640 simulations in total).

| **Parameter** | **Base value** | **SA** |
| --- | --- | --- |
| *Life history:* | | |
| average number of offspring ($b$) | 6 | 6 |
| maximum age ($age_m$) | 2 | 2 |
| number of breeding cycles in a year ($n_c$ ) | 6 | 6 |
| mate-search and dispersal distance ($D$) | 1, 2, 3 | $U_d${1, 2, 3} |
| probability of survival ($\omega$) | 0.53 | 0.53 |
| carrying capacity per patch ($K$) | 30 | 30 |
| female probability of mating ($p_r$) | 0.8, 1.0 | $U$[0.8, 1.0] |
| probability of polyandry ($p_m$) | 0.46, 0.8 | $U$[0.4, 0.7] |
| number of males mated per breeding cycle ($n_m$) | 2 | 2 |
| *Lethal* control*:* | | |
| reduction in survival with lethal control ($c_s$) | 0.15, 0.20 | $U$[0.1, 0.3] |
| *Release effort:* | | |
| number of introduction sites ($n_p$) | 16, 36, 64 | $U_d${16, 36, 64} |
| number of transgenic males introduced per site ($N_i$) | 10, 15, 20 | $U_d${5, 10, 15, 20} |
| number of releases over time ($n_i$) | 6,12 | $U_d${1, 2, …15} |

**Table S2.** The probability of eradication for each effort combination presented in Figure 2 under 'broad' ,'tactical' and 'tactical when detected' release strategies with gravid lethal, fsRIDL, and Y-linked editor (based on 720 simulations for each combination with various biological parameters; 116,640 simulations in total).

| Probability of eradication | Gravid lethal | | | fsRIDL | | | Y-linked editor | | |
|---|---|---|---|---|---|---|---|---|---|
| Strategy | Broad | Tactical | Tactical *w/* detection | Broad | Tactical | Tactical *w/* detection | Broad | Tactical | Tactical *w/* detection |
| 1 | 0.57 | 0.10 | 0.09 | 0.35 | 0.05 | 0.04 | 0.53 | 0.10 | 0.11 |
| 2 | 0.59 | 0.06 | 0.09 | 0.38 | 0.05 | 0.05 | 0.57 | 0.16 | 0.12 |
| 3 | 0.99 | 0.39 | 0.26 | 0.78 | 0.21 | 0.15 | 0.89 | 0.49 | 0.34 |
| 4 | **1.00** | 0.67 | 0.53 | 0.83 | 0.22 | 0.14 | 0.96 | 0.63 | 0.53 |
| 5 | **1.00** | 0.93 | 0.81 | 0.95 | 0.66 | 0.36 | 0.99 | 0.84 | 0.75 |
| 6 | **1.00** | **1.00** | **1.00** | 0.97 | 0.76 | 0.38 | **1.00** | 0.98 | 0.91 |
| 7 | 0.72 | 0.13 | 0.12 | 0.48 | 0.10 | 0.07 | 0.65 | 0.19 | 0.15 |
| 8 | 0.71 | 0.14 | 0.12 | 0.54 | 0.10 | 0.09 | 0.69 | 0.28 | 0.21 |
| 9 | **1.00** | 0.64 | 0.50 | 0.91 | 0.39 | 0.20 | 0.97 | 0.63 | 0.48 |
| 10 | **1.00** | 0.97 | 0.86 | 0.94 | 0.49 | 0.23 | **1.00** | 0.81 | 0.70 |
| 11 | **1.00** | 0.98 | 0.94 | 0.99 | 0.84 | 0.52 | **1.00** | 0.94 | 0.88 |
| 12 | **1.00** | **1.00** | **1.00** | **1.00** | 0.93 | 0.62 | **1.00** | **1.00** | 0.99 |
| 13 | 0.79 | 0.23 | 0.17 | 0.59 | 0.11 | 0.09 | 0.72 | 0.27 | 0.22 |
| 14 | 0.76 | 0.32 | 0.23 | 0.62 | 0.13 | 0.09 | 0.75 | 0.40 | 0.31 |
| 15 | **1.00** | 0.81 | 0.62 | 0.96 | 0.55 | 0.27 | 0.99 | 0.72 | 0.59 |
| 16 | **1.00** | **1.00** | 0.97 | 0.98 | 0.67 | 0.26 | **1.00** | 0.91 | 0.76 |
| 17 | **1.00** | **1.00** | 0.98 | **1.00** | 0.89 | 0.64 | **1.00** | 0.98 | 0.94 |
| 18 | **1.00** | **1.00** | **1.00** | **1.00** | 0.98 | 0.75 | **1.00** | **1.00** | **1.00** |

**Table S3.** The median expected time to eradication in years for each effort combination presented in Figure 2 under ‘broad‘ ,’tactical’ and ‘tactical when detected’ release strategies with Gravid Lethal, fsRIDL and Y-linked editor (based on 720 simulations for each combination with various biological parameters; 116,640 simulations in total).

| Time to eradication | | Gravid lethal | | | fsRIDL | | | Y-linked editor | | |
|---|---|---|---|---|---|---|---|---|---|---|
| | Strategy | Broad | Tactical | Tactical *w/* detection | Broad | Tactical | Tactical *w/* detection | Broad | Tactical | Tactical *w/* detection |
| | 1 | 3.50 | 1.67 | 1.33 | 3.50 | 2.00 | 1.50 | 3.50 | 2.67 | 2.17 |
| | 2 | 5.50 | 1.58 | 1.33 | 5.50 | 2.67 | 1.50 | 5.50 | 4.17 | 2.00 |
| | 3 | 3.67 | 3.00 | 1.67 | 3.67 | 3.00 | 1.67 | 3.67 | 3.33 | 2.50 |
| | 4 | 5.67 | 5.17 | 2.75 | 5.67 | 5.00 | 1.83 | 5.67 | 5.33 | 3.50 |
| | 5 | 3.83 | 3.50 | 2.00 | 3.83 | 3.67 | 2.00 | 3.83 | 3.67 | 2.83 |
| | 6 | 5.83 | 5.67 | 2.33 | 5.83 | 5.67 | 2.00 | 5.83 | 5.67 | 3.17 |
| | 7 | 3.50 | 2.83 | 1.33 | 3.50 | 2.83 | 1.67 | 3.50 | 3.25 | 2.33 |
| | 8 | 5.50 | 3.33 | 1.50 | 5.50 | 4.33 | 1.67 | 5.50 | 5.17 | 2.83 |
| | 9 | 3.67 | 3.33 | 1.83 | 3.67 | 3.50 | 1.83 | 3.67 | 3.50 | 2.83 |
| | 10 | 5.67 | 5.50 | 3.83 | 5.67 | 5.50 | 1.83 | 5.67 | 5.67 | 3.50 |
| | 11 | 3.83 | 3.83 | 2.00 | 3.83 | 3.83 | 2.17 | 3.83 | 3.83 | 2.83 |
| | 12 | 5.83 | 5.83 | 2.00 | 5.83 | 5.83 | 2.33 | 5.83 | 5.83 | 3.17 |
| | 13 | 3.67 | 3.00 | 1.42 | 3.67 | 3.00 | 1.67 | 3.67 | 3.33 | 2.50 |
| | 14 | 5.50 | 5.17 | 1.83 | 5.50 | 5.00 | 1.50 | 5.50 | 5.33 | 3.67 |
| | 15 | 3.83 | 3.50 | 2.00 | 3.83 | 3.67 | 2.00 | 3.83 | 3.67 | 2.83 |
| | 16 | 5.83 | 5.67 | 2.83 | 5.83 | 5.67 | 2.00 | 5.83 | 5.67 | 3.17 |
| | 17 | 3.83 | 3.83 | 2.00 | 3.83 | 3.83 | 2.33 | 3.83 | 3.83 | 2.83 |
| | 18 | 5.83 | 5.83 | 2.00 | 5.83 | 5.83 | 2.67 | 5.83 | 5.83 | 3.00 |

**Table S4.** The median number of transgenic individuals introduced for each effort combination presented in Figure 2 under ‘broad‘ ,’tactical’ and ‘tactical when detected’ release strategies with gravid lethal, fsRIDL and Y-linked editor (based on 720 simulations for each combination with various biological parameters; 116,640 simulations in total).

| Total inoculation effort | | Gravid lethal | | | fsRIDL | | | Y-linked editor | | |
|---|---|---|---|---|---|---|---|---|---|---|
| | Strategy | Broad | Tactical | Tactical *w/* detection | Broad | Tactical | Tactical *w/* detection | Broad | Tactical | Tactical *w/* detection |
| | 1 | 960.00 | 70.00 | 60.00 | 960.00 | 80.00 | 60.00 | 960.00 | 130.00 | 70.00 |
| | 2 | 1920.00 | 75.00 | 60.00 | 1920.00 | 100.00 | 50.00 | 1920.00 | 240.00 | 70.00 |
| | 3 | 2160.00 | 290.00 | 120.00 | 2160.00 | 260.00 | 115.00 | 2160.00 | 580.00 | 180.00 |
| | 4 | 4320.00 | 1150.00 | 310.00 | 4320.00 | 500.00 | 120.00 | 4320.00 | 1695.00 | 480.00 |
| | 5 | 3840.00 | 1235.00 | 280.00 | 3840.00 | 1400.00 | 220.00 | 3840.00 | 1715.00 | 450.00 |
| | 6 | 7680.00 | 3730.00 | 410.00 | 7680.00 | 4340.00 | 230.00 | 7680.00 | 4740.00 | 650.00 |
| | 7 | 1440.00 | 180.00 | 90.00 | 1440.00 | 180.00 | 90.00 | 1440.00 | 322.50 | 127.50 |
| | 8 | 2880.00 | 210.00 | 90.00 | 2880.00 | 315.00 | 90.00 | 2880.00 | 1080.00 | 165.00 |
| | 9 | 3240.00 | 825.00 | 217.50 | 3240.00 | 870.00 | 180.00 | 3240.00 | 1245.00 | 330.00 |
| | 10 | 6480.00 | 2505.00 | 645.00 | 6480.00 | 2790.00 | 180.00 | 6480.00 | 3397.50 | 630.00 |
| | 11 | 5760.00 | 2790.00 | 510.00 | 5760.00 | 3150.00 | 375.00 | 5760.00 | 3390.00 | 705.00 |
| | 12 | 11520.00 | 7987.50 | 555.00 | 11520.00 | 8512.50 | 420.00 | 11520.00 | 8760.00 | 915.00 |
| | 13 | 1920.00 | 340.00 | 120.00 | 1920.00 | 350.00 | 120.00 | 1920.00 | 580.00 | 180.00 |
| | 14 | 3840.00 | 860.00 | 160.00 | 3840.00 | 620.00 | 120.00 | 3840.00 | 1580.00 | 540.00 |
| | 15 | 4320.00 | 1420.00 | 370.00 | 4320.00 | 1680.00 | 240.00 | 4320.00 | 2100.00 | 500.00 |
| | 16 | 8640.00 | 4380.00 | 560.00 | 8640.00 | 5260.00 | 260.00 | 8640.00 | 5400.00 | 760.00 |
| | 17 | 7680.00 | 4520.00 | 660.00 | 7680.00 | 4880.00 | 560.00 | 7680.00 | 5060.00 | 990.00 |
| | 18 | 15360.00 | 11880.00 | 680.00 | 15360.00 | 12400.00 | 700.00 | 15360.00 | 12540.00 | 1080.00 |

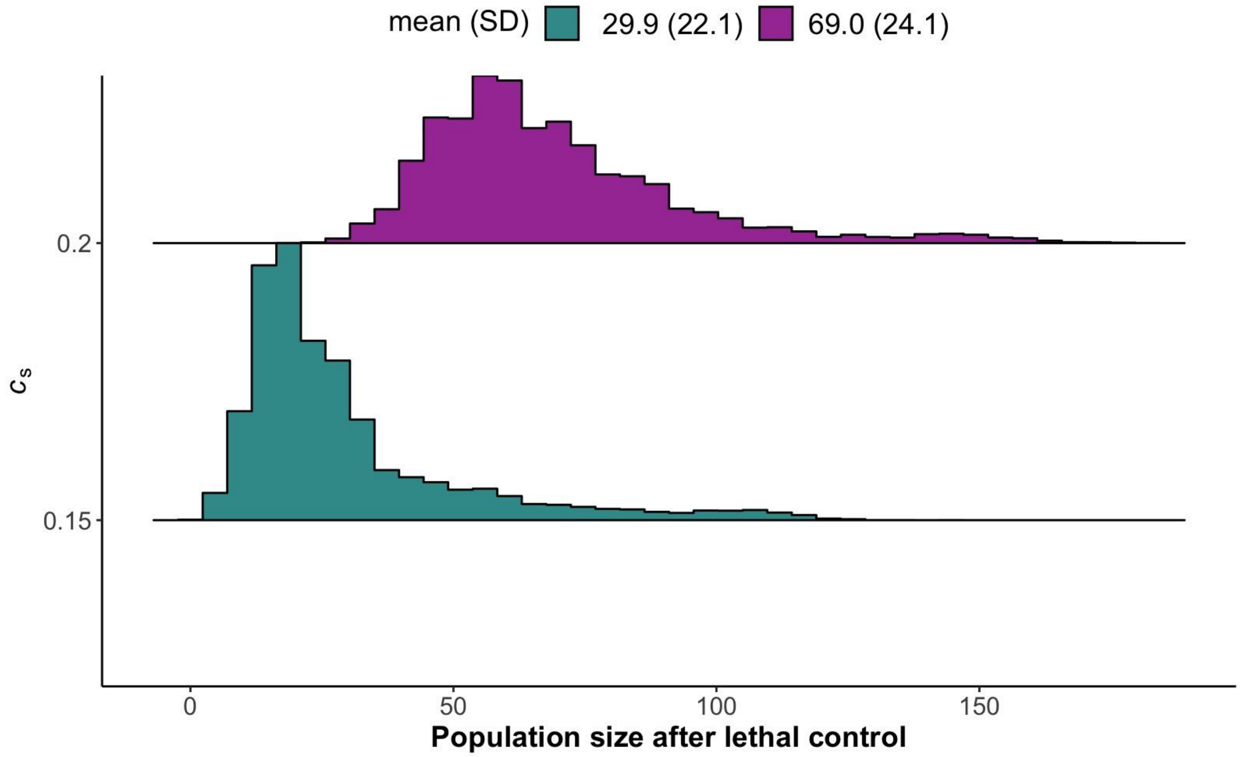


**Figure S1.** The distribution of population sizes remaining at the end of the lethal control program during which the survival probability was reduced by the multiplier $c_s$. (Based on 38880 simulations; in 2 simulations lethal control (with $c_s$ = 0.15) resulted in eradication before introduction of transgenic individuals, hence are excluded from further analyses.)

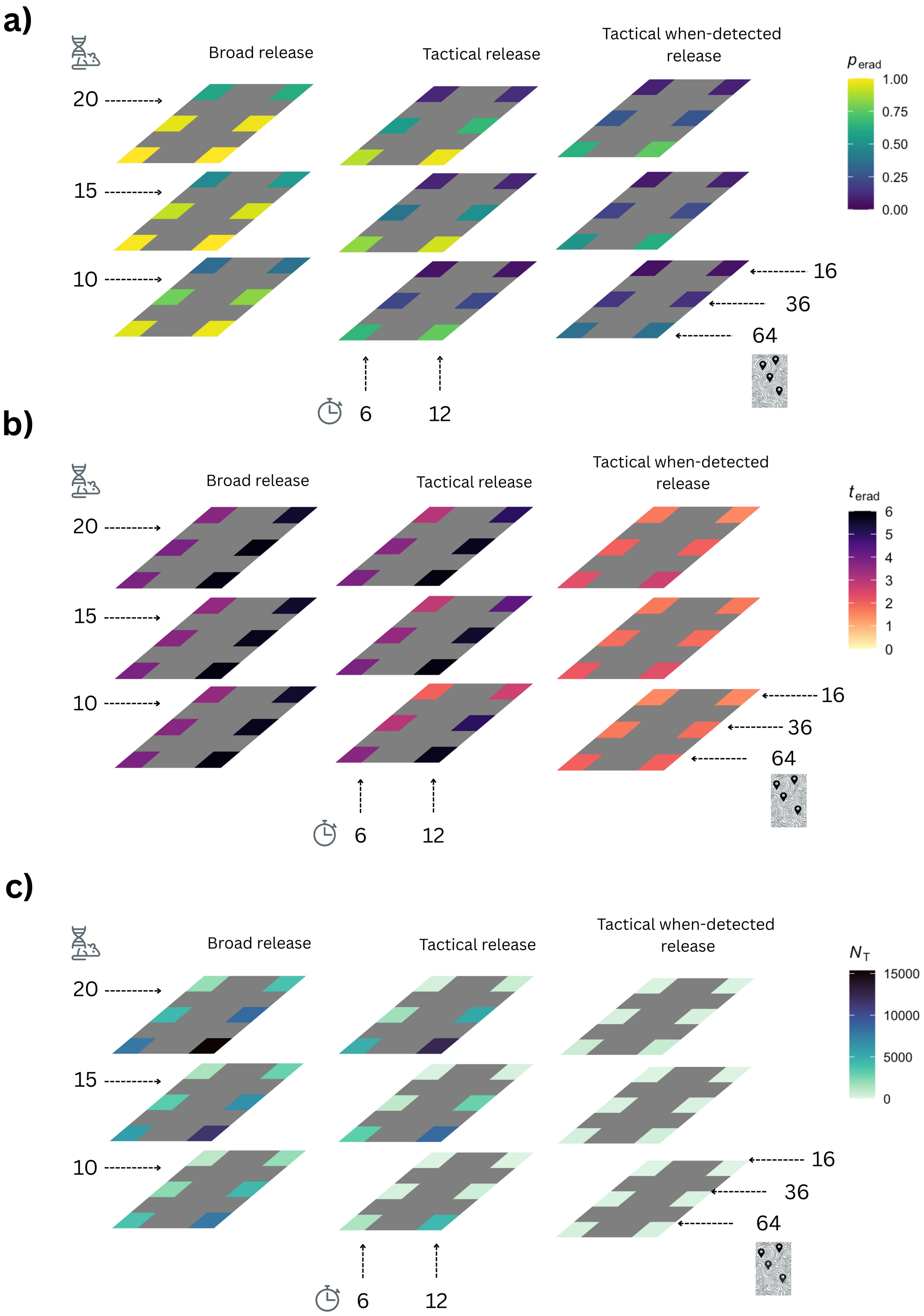
a)
Broad release
Tactical release
Tactical when-detected release
$p_{erad}$
1.00
0.75
0.50
0.25
0.00
20
15
10
16
36
64
6
12
b)
Broad release
Tactical release
Tactical when-detected release
$t_{erad}$
6
5
4
3
2
1
0
20
15
10
16
36
64
6
12
c)
Broad release
Tactical release
Tactical when-detected release
$N_T$
15000
10000
5000
0
20
15
10
16
36
64
6
12

**Figure S2.** fsRIDL **a)**. The probability of eradication for each release effort combination under ‘broad’, ‘tactical’, and ‘tactical when-detected’ release strategies. **b)** The median time to eradication (in years) for simulations when eradications were successful. **c)** The median number of transgenic individuals introduced for each effort combination. (See the caption of Figure 3 in the main text for more details).

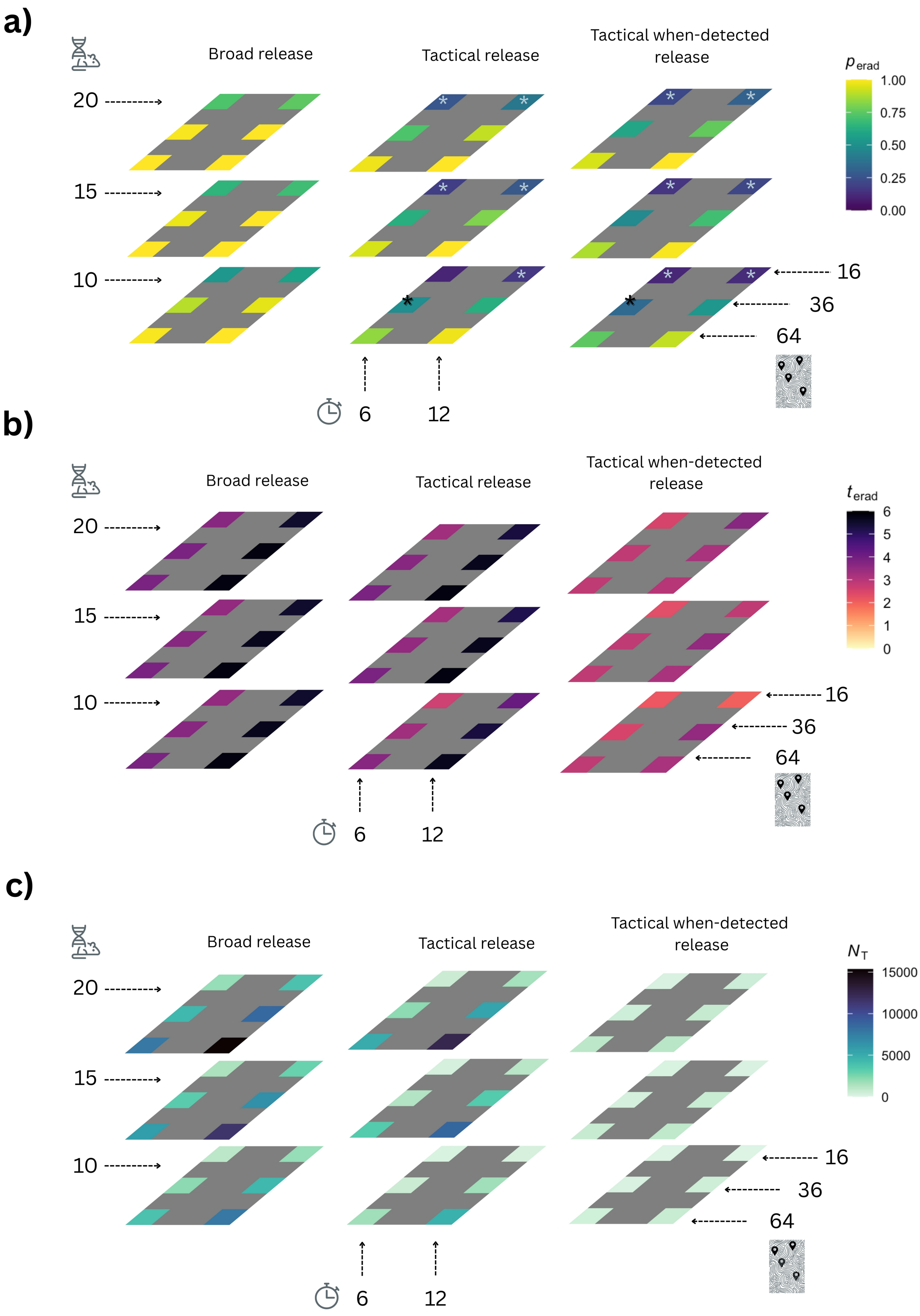

a)
Broad release
Tactical release
Tactical when-detected release
$p_{erad}$
1.00
0.75
0.50
0.25
0.00
20
15
10
16
36
64
6
12
b)
Broad release
Tactical release
Tactical when-detected release
$t_{erad}$
6
5
4
3
2
1
0
20
15
10
16
36
64
6
12
c)
Broad release
Tactical release
Tactical when-detected release
$N_T$
15000
10000
5000
0
20
15
10
16
36
64
6
12

**Figure S3.** Y-linked editor **a)**. The probability of eradication for each release effort combination under ‘broad’, ‘tactical’, and ‘tactical when-detected’ release strategies. The release efforts where Y-linked editor outperformed Gravid Lethal are displayed with ‘*’. **b)** The median time to eradication (in years) for simulations when eradications were successful. **c)** The median number of transgenic individuals introduced for each effort combination. (See the caption of Figure 3 in the main text for more details).

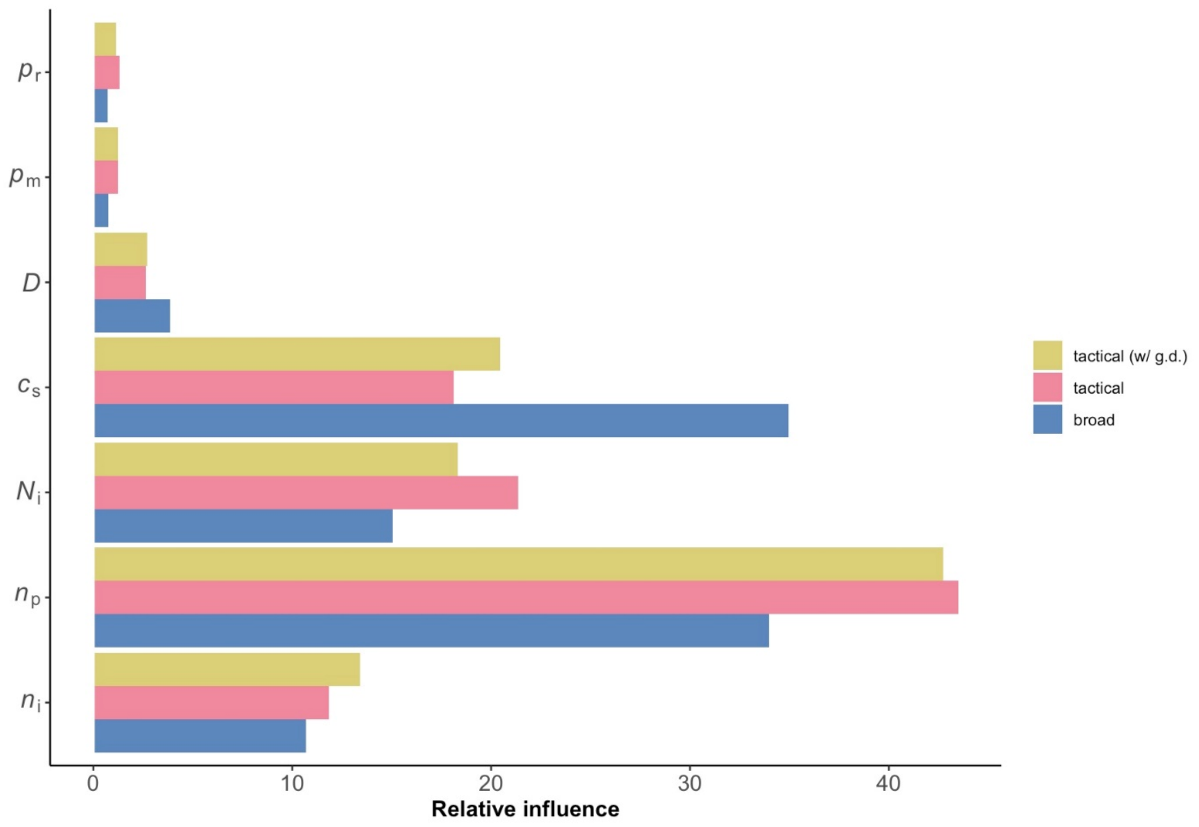


**Figure S4.** The relative influence of parameters on the probability of eradication with the gravid lethal genetic biocontrol (simulations in which eradication with classic control was successful before the application of genetic biocontrol are excluded), based on 10,000, 9946, and 9961 simulations using broad, tactical, and tactical when detected release strategies, respectively. Parameter abbreviations are provided in Table S1.
